# Combining experiments and simulations to examine the temperature-dependent behaviour of a disordered protein

**DOI:** 10.1101/2023.03.04.531094

**Authors:** Francesco Pesce, Kresten Lindorff-Larsen

**Affiliations:** Structural Biology and NMR Laboratory, The Linderstrøm-Lang Centre for Protein Science, Department of Biology, University of Copenhagen, Copenhagen, Denmark

## Abstract

Intrinsically disordered proteins are a class of proteins that lack stable folded conformations and instead adopt a range of conformations that determine their biochemical functions. The temperature-dependent behaviour of such disordered proteins is complex and can vary depending on the specific protein and environment. Here, we have used molecular dynamics simulations and previously published experimental data to investigate the temperature-dependent behaviour of Histatin 5, a 24-residue-long polypeptide. We examined the hypothesis that Histatin 5 undergoes a loss of polyproline II structure with increasing temperature, leading to more compact conformations. We found that the conformational ensembles generated by the simulations generally agree with small-angle X-ray scattering data for Histatin 5, but show some discrepancies with the hydrodynamic radius as probed by pulsed-field gradient nuclear magnetic resonance spectroscopy, and with the secondary structure information derived from circular dichroism. We attempted to reconcile these differences by reweighting the conformational ensembles against the scattering and NMR data. By doing so, we were in part able to capture the temperature-dependent behaviour of Histatin 5 and to link the observed decrease in hydrodynamic radius with increasing temperature to a loss of polyproline II structure. We were, however, unable to achieve agreement with both the scattering and NMR data within experimental errors. We discuss different possibilities for this outcome including inaccuracies in the force field, differences in conditions of the NMR and scattering experiments, and issues related to the calculation of the hydrodynamic radius from conformational ensembles. Our study highlights the importance of integrating multiple types of experimental data when modelling conformational ensembles of disordered proteins and how environmental factors such as the temperature influence them.

## Introduction

Intrinsically disordered proteins (IDPs) are a class of proteins characterized by the lack of stable folded conformations. Instead, IDPs may adopt a wide range of conformations, which help determine their biochemical functions.^1,2^ This ensemble of conformations, although disordered, may show sequence-dependent populations of specific regions of the Ramachandran map,^3^ and the formation of transient secondary structures is common.^4,5^ Therefore, they are better described as statistical coils.^6^ Biophysical measurements make it possible to get insights into the propensity of IDPs to form (transient) secondary structures, although the interpretation of ensemble-averaged data is often ambiguous.^7^ Alternatively, molecular dynamics (MD) simulations enable direct access to the population of conformations of IDPs at atomistic resolution, but its utility may be limited by difficulties in sampling and by the accuracy of the force fields used.^8–12^

The complexity of the problem increases even more when considering that the behaviour of IDPs is often highly dependent on the environment.^13^ One factor that may affect the behaviour of IDPs is the temperature.^14–23^ The temperature-dependence of the conformational properties of IDPs may, however, be complex, as they depend on the combined effects of many different physical effects including the hydrophobic effect, changes in local structure, and other entropic contributions, and may vary depending on the specific protein and the specific environment in which it is found.^24–26^ Other than influencing the single chain behaviour, temperature is also known to influence how IDPs interact and their tendency to form high-order entities.^27,28^ Therefore, understanding how temperature affects the single-chain behaviour, may provide new insights into how aggregates and biomolecular condensates can form as a consequence of particular single-chain behaviours. ^29^

Experimentally, the temperature-dependent behaviour of the local structure of IDPs has previously been studied using for example circular dichroism (CD) spectroscopy,^4,5, 30–32^ where changes in the spectra, although difficult to interpret at atomistic resolution, have been suggested to be due to formation of *α*-helices and/or loss of polyproline II (PPII) structures with increasing temperature.^15,23,33,34^ This change in propensity to form certain secondary structure elements has also been linked to changes in the overall dimension of IDPs.^5,14,15,23,35^ For example, it has been suggested that the loss of PPII with increasing temperature is responsible for the conformational ensemble of certain IDPs to get more compact.^15,23^ Even though this might sound counter intuitive, it could be explained by the fact that the PPII structure is a particularly extended secondary structure element, though other mechanisms are also likely to be important.^14,24,25^

The overall chain dimension of IDPs can be probed, for example, by small-angle X-ray scattering (SAXS) or pulsed-field gradient (PFG) NMR spectroscopy. The former yields structural information that are often linked to the average radius of gyration (*R*_g_) of an IDP in solution,^36^ while the latter yields the translational diffusion coefficient, which is in turn related to the average hydrodynamic radius (*R*_h_) of an IDP in solution.^37^ *R*_g_ and *R*_h_ are both measures that describe chain compaction, and are therefore correlated.^38,39^ Nonetheless, SAXS and PFG NMR may sometimes be used to probe subtly different aspects of compaction.^38^ Specifically, SAXS probes the ensemble averaged 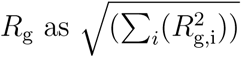 for each conformation *i* in the solution ensemble. Conversely, the translational diffusion coefficient measured by PFG NMR, when converted into an *R*_h_ is averaged as 1/∑*_i_*(1/*R*_h,i_). By consequence of this different averaging, compaction measured by SAXS may put greater weight on expanded conformations, while compaction measured by PFG NMR is more representative of compact conformations.^38^ Both theory and numerical calculations show that the*R*_g_*/R*_h_ depends on compaction^38–40^ and so a joint analysis of SAXS and PFG NMR measurements may provide additional information compared to analysing these individually.^38,41^

Here, we use the small protein Histatin 5 (hereafter Hst5) to study the relationship between local and global structure of a disordered protein and the effect of temperature on these properties. This 24 residue-long polypeptide has previously been characterized by a multitude of experiments.^23,42–47^ CD spectra recorded at different temperatures between 283 K and 323 K suggested that a loss of PPII structure takes place with increasing temperature. Increasing the temperature also results in a decrease of the *R*_h_ measured by PFG NMR, while the *R*_g_ extracted from SAXS data appears not to be particularly sensitive to temperature. Previous attempts of computational characterization of the conformational ensembles of Hst5 at different temperatures did not completely succeed in explaining the experimental results,^23^ but it was suggested that the drop in *R*_h_ with increasing temperature might be due to the temperature-dependent change in the propensity of forming PPII structures.

To provide a structural explanation of the experimental data, we extensively sample the conformational ensemble of Hst5 at different temperatures by running MD simulations enhanced with metadynamics.^48^ We then compare the conformational ensembles to the available temperature-dependent experimental data, but do not observe substantial temperature-dependent changes in compaction and propensity to form PPII structures. Inspired by previous work that combined experiments and simulations to reconcile different experimental observations,^49–51^ we use the Bayesian/Maximum-entropy reweighting approach^52–54^ to refine the simulations against SAXS and *R*_h_ from PFG NMR.^41^ We find that when we improve the agreement with the *R*_h_ we also observe a temperature-dependent change in the amount of PPII structure in qualitative agreement with the CD data, therefore providing a molecular-level description of a link between chain compaction and propensity to form PPII-like structure. Conversely, when we combine the simulations with the SAXS data we do not observe any temperature-dependent change in compaction and PPII structure content. When we refine the ensembles simultaneously against the SAXS and NMR data, we were not able to achieve the same level of agreement as when we optimized against them individually. We discuss possible origins of this observation, including minor differences in the conditions of the experiments, force fields inaccuracies, and issues related to how *R*_h_ values are estimated from conformational ensembles of IDPs.

## Results and Discussion

### Comparison of MD simulations and experiments

We sampled the conformational ensembles of Hst5 at 283 K, 293 K, 310 K and 323 K with MD simulations using the all-atom Amber ff99SB-disp (a99SB-disp) force field,^11^ as this has previously been shown to be reliable in capturing the overall chain dimension of IDPs^11^ and in reproducing NMR scalar couplings of a peptide with a high PPII structure propensity. ^55^ In order to improve the sampling of the overall dimensions and secondary structure preferences of Hst5, we used Parallel Bias Metadynamics^56^ (PBMetaD; see Methods) simulations with two semi-independent replicas biasing the same free-energy surface (FES).^57^ We ran these simulations until the sampling of the collective variables (CVs) that we biased with PBMetaD converged within a reasonable error (*i.e.* an average error on the estimated free-energy surface (FES) of the *R*_g_ lower than 1 kJ/mol; Fig. S1). We achieved this in all cases in a simulation time comprised between 2 and 2.5 *µ*s per replica. We also verify that, along the simulations, both *R*_g_ and the backbone dihedrals diffuse in the CV space and do not get stuck in a specific state, supporting that the extent of biasing used in the simulations is sufficient for the system to escape local minima (Fig. S2 and S3). Furthermore, because of the well-tempered biasing^58^ (see Methods), it would be ideal for the amount of bias dispensed in the system, which decreases over time, to reach a quasi-static regime, to ensure a proper convergence of the simulation. This happens when all accessible regions of the FES have been explored. In our simulations, we observe that overall the bias deposition and free-energy surfaces become quasi-static over time (Fig. S4), with the exception when sampling frames whose *R*_g_ is outside the interval in which the bias is active (Fig. S4b). Therefore these few frames are not taken into account when calculating averages and errors (see Methods).

We then calculated SAXS intensities, *R*_g_, *R*_h_, and per-residue secondary structure propensities, and compared these to the experimental values. We find that all simulations—apart from that at 310 K—provide a *χ*^2^ between the calculated and experimental SAXS data close to 1 (Fig. 1a), which, given the use of normalized error bars for SAXS intensities (see Methods), shows that the ensembles accurately fit the SAXS experiments. As previously reported,^23^ we find that the *R*_g_ values derived from the experimental SAXS profile (with the Guinier approximation, using the ATSAS package^59^) show that the average dimension for Hst5 in solution at the different temperatures are very similar (Fig. 1b). Similarly, the conformational ensembles sampled by MD provide average *R*_g_ values that are all comparable when taking the errors into account (with a subtle trend of expansion with increasing temperature), but consistently smaller than those from the Guinier approximation. This observation is in line with the expectation that SAXS-derived *R*_g_ carries the contribution of the protein hydration layer,^60^ while from the conformational ensembles the *R*_g_ is calculated on the coordinates of the protein atoms only.^61^ For this reason we only rely on the *χ*^2^_*r*_ to SAXS data to determine whether a certain ensemble is accurate in terms of chain dimensions as described by SAXS data, since in calculating the SAXS we take into account both the protein hydration layer and the excluded volume effect (see Methods).

**Figure 1:**
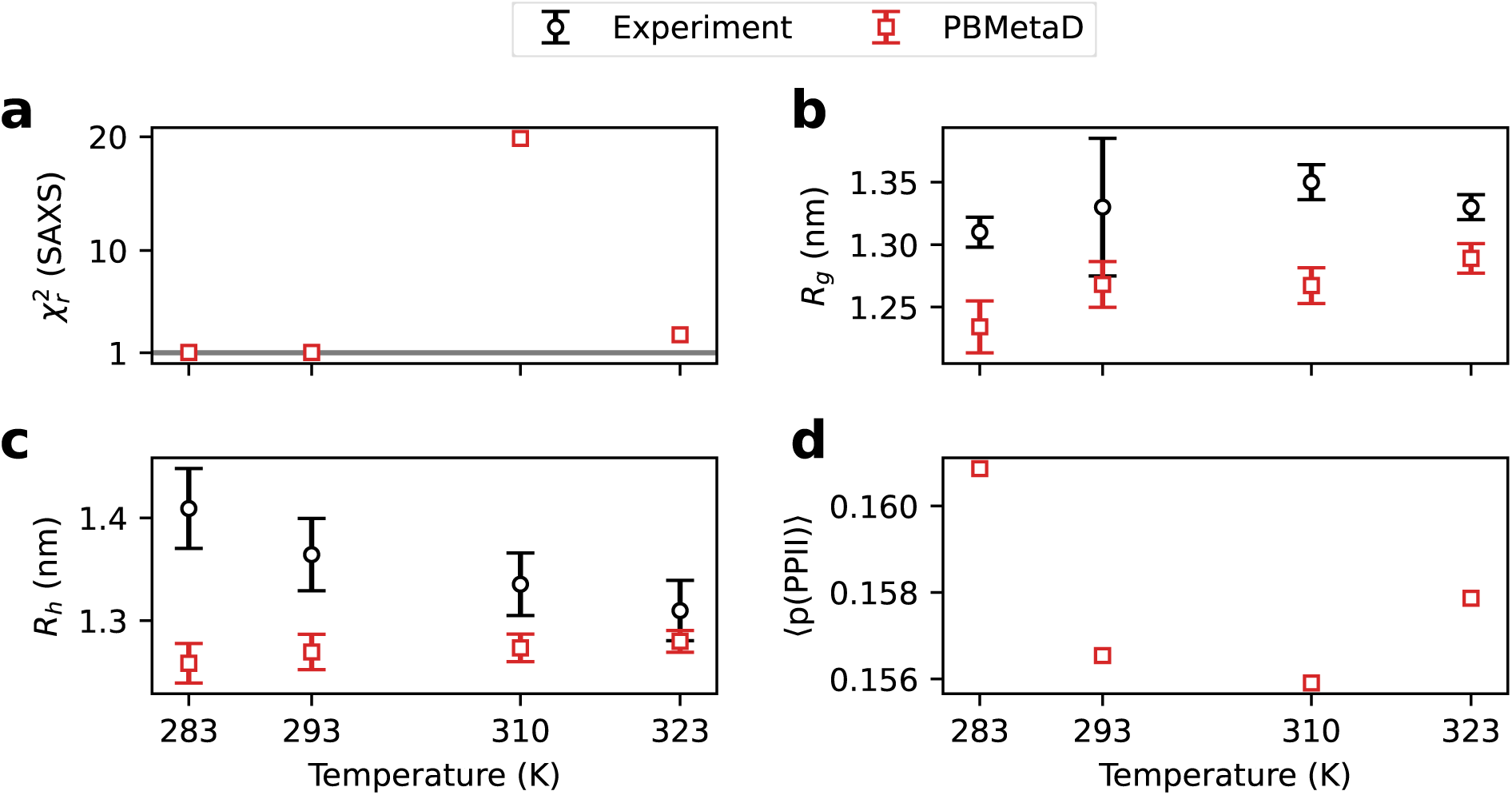
Average structural features calculated from MD simulations (red squares) at four different temperatures are compared to their experimental counterparts (black circles, when applicable). (a) *χ*^2^ to SAXS data (grey horizontal line indicates *χ*^2^_*r*_=1, that denotes ensembles that correctly fit the experiments; see Methods), (b) *R*_g_, (c) *R*_h_, (d) PPII structure content. As discussed in the main text, the deviation in *R*_g_ (panel b), despite good agreement with the SAXS intensities (panel a), may likely in part be attributed to the contribution of the solvation layer to the *R*_g_ derived from Guinier fitting. ^61^

The calculated *R*_h_ values appear to be consistently smaller than the experimental values determined by PFG NMR at any of the temperatures (Fig. 1c). This observation might indicate that the a99SB-disp force field underestimates the *R*_h_ of Hst5. However, we note a number of complications when calculating the *R*_h_ from conformational ensembles of IDPs^62,63^ as well as issues related to how the PFG NMR measurements are used to estimate *R*_h_.^63^ Specifically, we recently showed^63^ that the *R*_h_ calculated with HullRadSAS^64^ provides the best overall agreement with experiments across a series of IDPs. Nonetheless, we note that this observation depends on the *R*_h_ value used for the reference compound (here 1,4-dioxane^63,65^) that is used to derive the *R*_h_ from the diffusion data.^62,63^ Thus, earlier values of the *R*_h_ of dioxane^65^ suggested that a different approach to calculate *R*_h_ was more accurate.^62^ Finally, the comparison between averages of *R*_h_ values calculated for individual conformations and the translational diffusion coefficient rests on the assumption of treating *R*_h_ as an ensemble- and time-averaged quantity.^66^ Together, these issues mean that it is difficult to determine the origin of the deviation between experiments and simulations.

The CD spectra of Hst5 show profiles often taken to represent PPII structure. The decrease in intensity of the PPII signature with increasing temperature suggests that this secondary structure element is less populated with increasing temperature.^23^ Comparing CD spectra to simulations is difficult as previous attempts at calculating CD spectra from conformational ensembles of disordered peptides have shown large discrepancies between the experimental and calculated spectra.^55,67,68^ Also, we are not aware of approaches to deconvolute CD spectra of disordered proteins that takes into account PPII-like signals and which can accurately extract the average content of PPII structure. For these reasons, we instead perform a qualitative comparison of our ensembles to the CD data of Hst5 by calculating the average content of PPII structure in our ensembles and examine its temperature dependence. We find that the ensembles do not show any substantial differences in PPII structure at the different temperatures, with changes of at most 0.6% (Fig. 1d). All together these observations suggest that the force field underestimates the *R*_h_ from PFG NMR and does not show changes in compaction and propensity to form PPII structure with temperature.

### Refinement of the simulations against SAXS experiments

To test the hypothesis that a decrease in chain dimension comes with a lowered propensity to form PPII structure, we improved the agreement between the ensembles and data that encode information about the protein dimension at different temperatures. We first used the iBME^69^ approach to reweight the ensembles against SAXS data; in this way we construct conformational ensembles that are in better agreement with experiments, but with minimal perturbations to the ensembles generated by the force field alone. With very mild reweighting (Fig. S5), we achieved a good agreement with SAXS data for all simulations (Fig. 2a,b), but without any major change in *R*_h_ (Fig. 2c) and PPII structure propensity (Fig. 2d), and only the ensemble simulated at 310 K getting slightly more compact as seen by both *R*_h_ and *R*_g_ (Fig. 2c and S6). We note that the larger *χ*^2^_*r*_ before reweighting for the simulation at 310 K (and to a minor extent also that at 323 K) is likely due to a higher signal-to-noise ratio in the experimental SAXS profile at these temperatures compared to those at 283 K and 293 K (Fig. 2a). This difference arises from the use of more concentrated samples in the SAXS measurements at 310 K and 323 K.^23^ This, and the fact that we do not observe major changes in the ensembles upon reweighting against SAXS, suggest that the ensembles from PBMetaD fit the experimental SAXS data well at all temperatures and differences in ^2^ are due to different magnitudes of the error bars among the experiments at different temperatures. Indeed the normalized average error bars (see Methods) are 20%, 14%, 3% and 8% for SAXS data collected respectively at 283 K, 293 K, 310 K and 323 K. This suggests that the differences in *χ*^2^_*r*_ observed are in part due to the different magnitude of the errors across the experiments, though issues related to convergence and/or force field inaccuracy might also be in play. As seen also from the *R*_g_ values extracted from the experimental SAXS profiles, the SAXS data do not suggest any substantial change in average chain dimension with temperature. Nonetheless, even though the SAXS data does not change substantially as a function of temperature, this does not mean that the temperature may not influence the average dimension of Hst5 in solution, as SAXS experiments only probe certain aspects of chain compaction.^38,49,50,62^ We note also the difference between the *R*_g_ value calculated from the protein coordinates and from a Guinier analysis of SAXS data calculated from the same coordinates (Table S6), and that such differences have previously been shown to arise from a contribution from the solvent to the SAXS data.^61^

**Figure 2:**
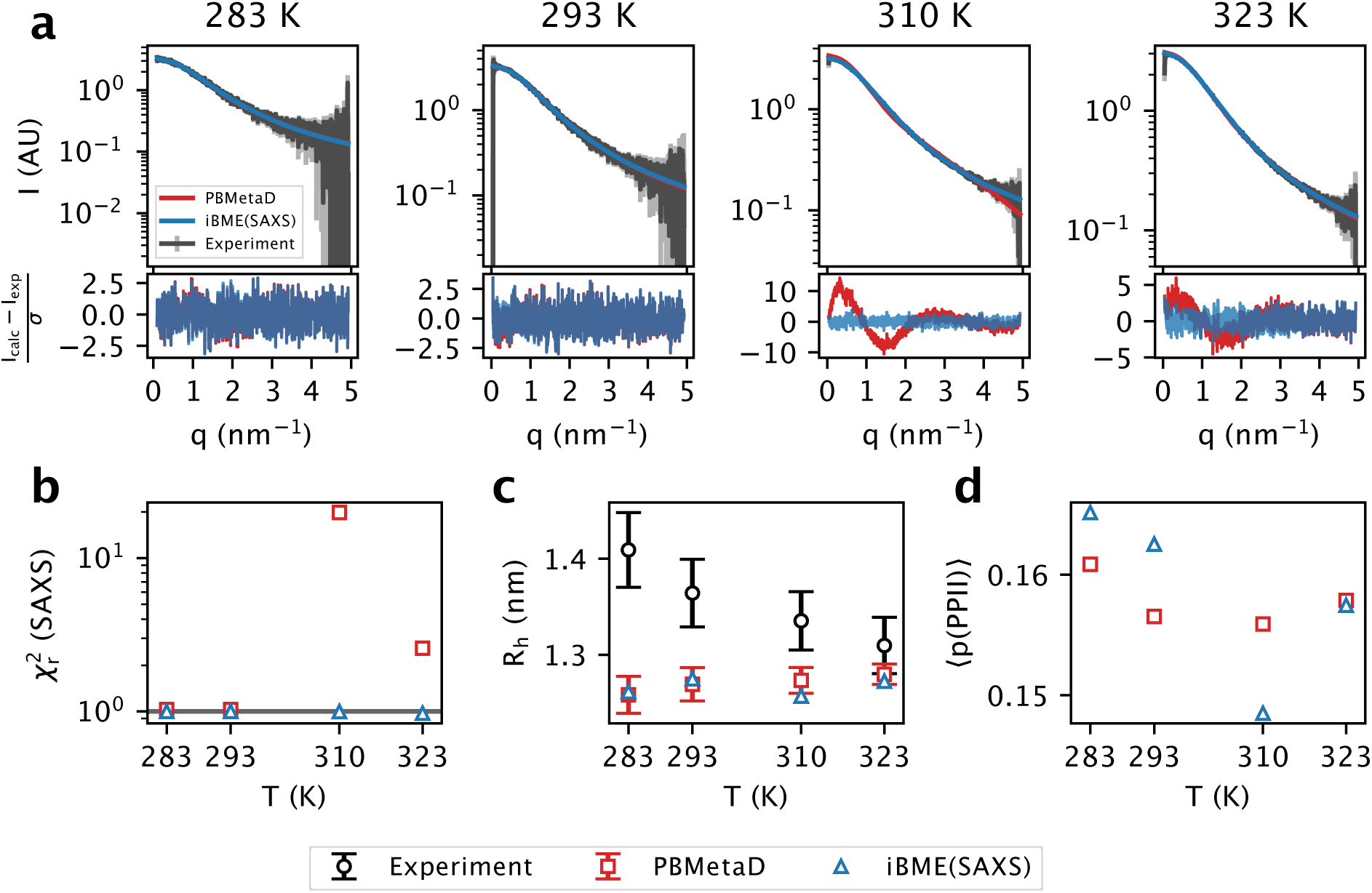
Refinement of the simulations against SAXS data. (a) SAXS profiles of prior (red) and reweighted (blue) ensembles compared with the experiment (grey). (b) *χ*^2^_*r*_ calculated from the profiles in panel a. (c) Ensemble-averaged *R*_h_ calculated with either the prior or SAXS-refined weights. (d) Ensemble-averaged per-residue PPII structure population calculated with either the prior or SAXS-refined weights.

### Refinement of the simulations against ***R***_h_ from PFG NMR

We also reweighted the simulations against the *R*_h_ from PFG NMR independently from the SAXS data. With the level of reweighting that we applied (Fig. S7), the reweighted *R*_h_ values from the simulations at 293 K, 310 K and 323 K come within the error bars associated to the experimental values, while the reweighted *R*_h_ of the simulations at 283 K remains below the experimental value (Fig 3a). The sampling at 283 K is more restricted to compact conformations and lacks expanded conformations whose population could be increased by reweighting (Fig. S8). Indeed, by reweighting the simulation at 323 K against the experimental *R*_h_ recorded at the lower temperatures, in all cases (but in particular for the lowest temperature) we achieve a better agreement with the experiments (Fig. S9), because there are more expanded conformations that can be up-weighted to fit the experiments (Fig. S8). Remarkably, when improving the agreement with the experimental *R*_h_, we also observe an overall increase in the amount of PPII structure, that follows the same trend suggested by CD data (Fig. 3b). Observing the relationship between conformational *R*_h_ and number of residues in PPII conformation, we see that, while conformations with only few residues involved in a PPII structure can be either very compact or very expanded, conformations with many residues in a PPII structure are all expanded (Fig. 3c). Indeed, by reweighting the ensembles against the experimental *R*_h_ values (that likely increases the weight of conformations with larger *R*_h_), we also observe that conformations with five to ten residues involved in a PPII structure are up-weighted, while conformations with no or little of this secondary structure element are down-weighted (Fig. 3d). The down-side arising from reweighting the simulations of Hst5 against *R*_h_ is that the agreement with SAXS data decreases (Fig. 3e), that also translates in the ensemble-averaged *R*_g_ becoming larger (Fig. S6).

**Figure 3:**
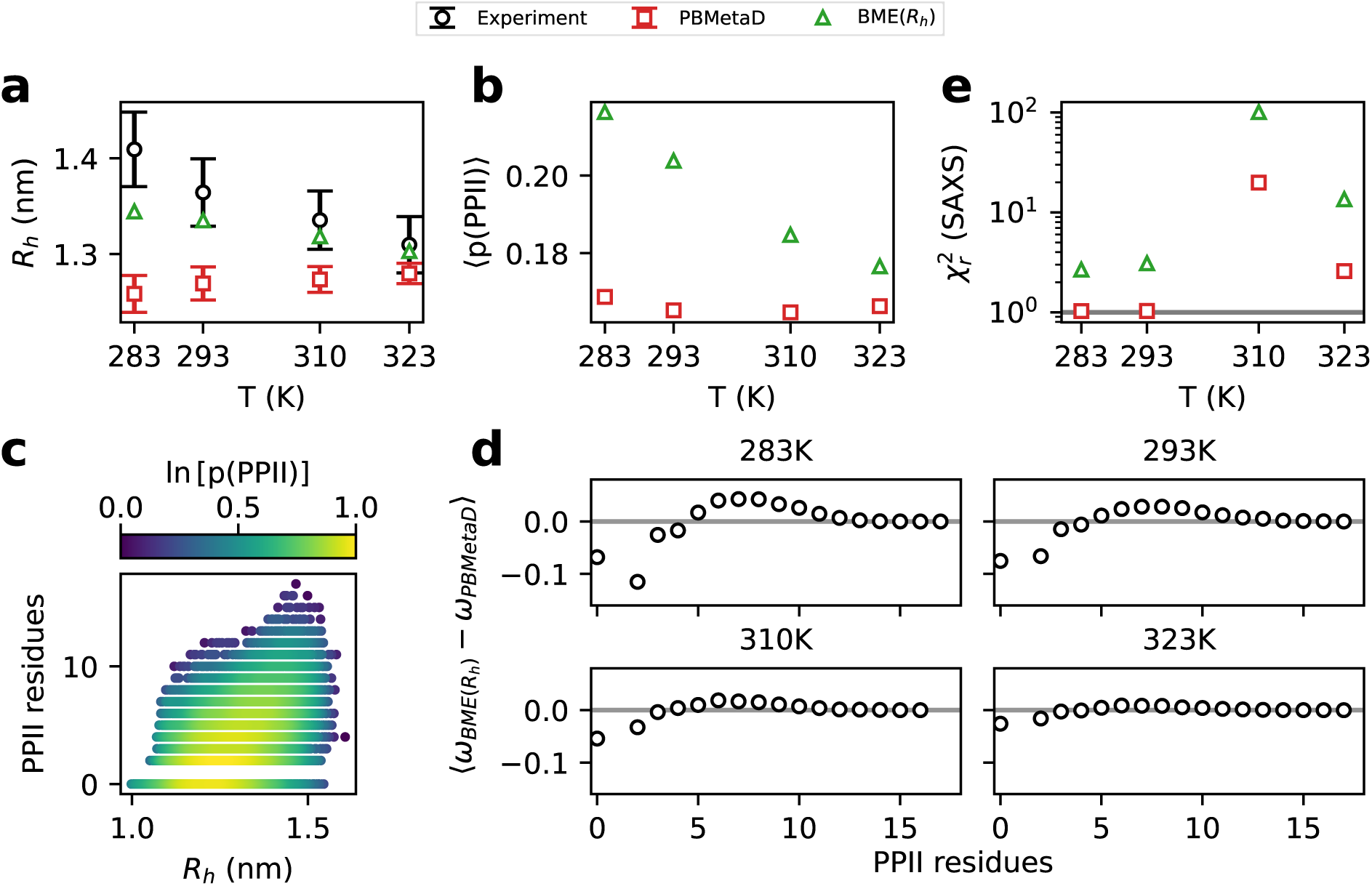
Refinement of the simulations against *R*_h_ from PFG NMR measurements. We show the (a) *R*_h_, (b) average PPII structure propensity and (e) *χ*^2^_*r*_ to SAXS data for the expectation value from the experiment (black, when applicable), the ensemble average from the simulations (red) and the simulations reweighted against the experimental *R*_h_ (green). (c) Density plot showing the number of residues involved in a PPII structure as a function of the conformational *R*_h_, together with their relative frequency. (d) Average difference between *R*_h_-refined weights and prior weights, for frames with a certain number of residues involved in forming a PPII structure.

### Simultaneous refinement of the simulations against experimental *R***_h_ and SAXS**

As discussed above, SAXS and *R*_h_ can probe different aspects of the compaction of protein conformations; therefore refining against both experiments might in principle provide additional information compared to analysing them individually. We therefore refined the ensembles simultaneously against SAXS and *R*_h_ (see Methods). We note here one complication when refining simultaneously against different sources of experiments. Specifically, setting the appropriate balance between the force field and the experiments depends (among other things) on both the experimental errors and the errors associated with how we calculate observables from experiments.^52,54,70^ Thus, when setting the balance between SAXS and the PFG NMR data we are assuming that the relative balance between these experiments is correct as the reweighting approach only tunes the balance between the simulations and the experiments via a single parameter.

With these caveats in mind, we find that the results of the combined reweighting (Fig. 4) are similar to that obtained when reweighting against SAXS alone (Fig. 2). The main reason for this is that in this combined reweighting the SAXS experiments overall get a substantially greater weight due to a combination of the larger number of data points and the associated errors. Specifically, the SAXS experiments are represented by a large number of intensity measurements while the experimental *R*_h_ is represented by a single data point (that itself is derived from a small number of points in the PFG NMR experiments). Therefore, if the reweighting cannot satisfy both experiments simultaneously, it will tend to optimize mostly on the experiment with the largest number of data points, that is the SAXS experiments, unless the other experiments have much smaller errors. To demonstrate this effect, we changed the balance of the reweighting by computationally expanding the number of data points for the *R*_h_ to the same number of data points available in the SAXS profile. When reweighting against this alternative dataset we find a solution that shows intermediate level agreement to both sets of experiments (Fig. S10).

**Figure 4:**
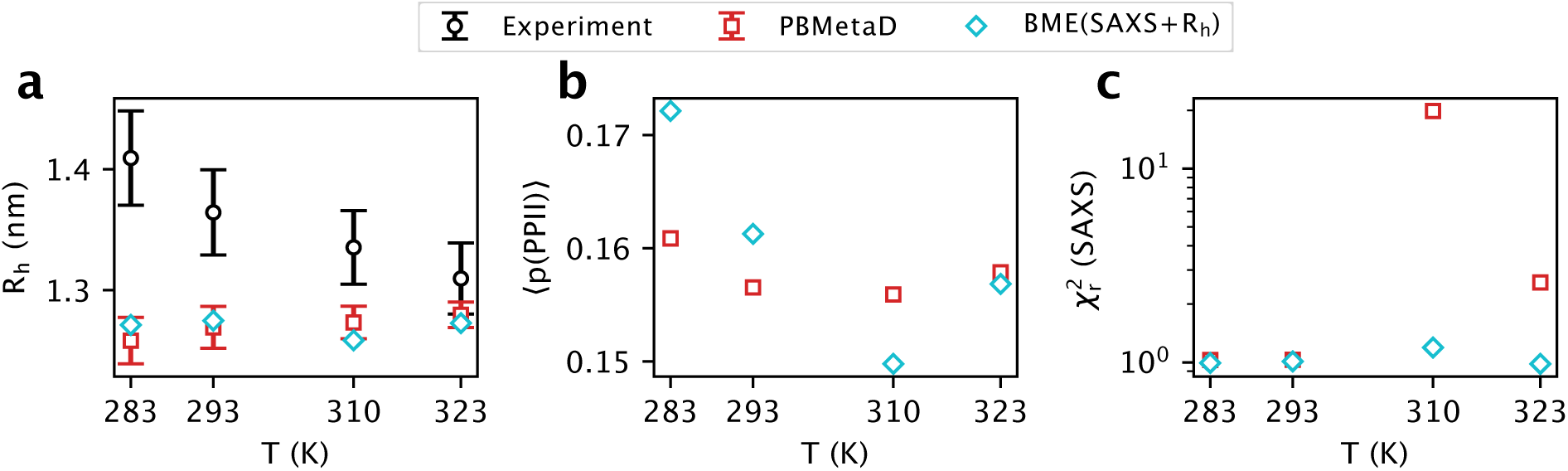
Refinement of the simulations against SAXS and *R*_h_ simultaneously. We show the (a) *χ*^2^_*r*_ to SAXS data (b) *R*_h_ and (c) average PPII structure propensity from the experiment (black, when applicable), the ensemble average from the simulations (red) and the simulations reweighted against both SAXS and *R*_h_ (cyan).

## Conclusions

Integrative modelling makes it possible to provide atomistic models of IDPs by combining information from experiments probing ensemble-averaged structural features with conformational models generated for example by MD simulations. Taken alone, many experiments only provide information about a specific structural feature, for example the average chain dimension, while being mostly insensitive to other properties of the conformational ensemble. Therefore, integrating simultaneously multiple sources of independent experimental information can provide a more realistic determination of the conformational ensemble of an IDP.^6,38,41,49,50,71^

We have here focused on Hst5 as a case study, in an attempt to explain the previously observed trend showing the *R*_h_ from PFG NMR decreasing with temperature, together with a drop of the PPII structure signal from CD spectra^23^ and a relatively temperatureindependent *R*_g_ from SAXS. While Hst5 is relatively simple in terms of its conformational preferences, its small size makes it useful because it is easier to sample by means of allatom molecular simulations. This in turn makes it possible to study small differences across different conditions as well as potential coupling between local and global conformational preferences. We find that the simulations alone were mostly in good agreement with the SAXS data, but did not appear to reproduce the *R*_h_ from PFG NMR and the loss of PPII structure with increasing temperature. We therefore applied a reweighting approach to refine the ensembles to achieve a better agreement with the experiments. While we could refine against the SAXS and NMR data separately, we were not able to achieve simultaneous agreement with the SAXS and *R*_h_ data within experimental errors. Refining against the CD data would have been interesting, but is hampered by lack of methods to calculate accurate CD spectra for disordered proteins; however, we find that refining against the *R*_h_ provided a picture in qualitative agreement with the previous interpretation of the CD data.

In light of our observations, one conclusion is that while the force field used provides an overall good agreement with the SAXS data, it does not appear to capture the temperature-dependence shown by the experimental NMR and CD data. This force field, as other force fields shown to be relatively accurate for disordered proteins, has been tested to reproduce *R*_g_ extrapolated from SAXS data, and indeed it does a good job in reproducing that experimental information. When it comes to *R*_h_, it seems that the force field overall underestimates the *R*_h_, with the exception of that at 323 K. We note, however, that the observed disagreement between the simulations and the *R*_h_ values from PFG NMR may also be due to inaccuracies in the forward model for *R*_h_ ^62,63^ as well as remaining uncertainty about how to determine *R*_h_ experimentally.^63^ The possibility to calculate accurately CD spectra from conformational ensembles would also aid in understanding the reliability of the forward model for the *R*_h_. Given the observed correlation between *R*_h_ and PPII structure content, a quantitative comparison to CD spectra could provide direct information about the possibility that the force field underestimates the PPII structure content of Hst5, together with its *R*_h_. In absence of such a model, we decided to rely on a qualitative analysis of the secondary structures. Based on the backbone dihedral angles, we can calculate the per-residue secondary structure for each frame of our simulations. Doing so, we did not find any temperature dependence on the amount of PPII structure in our simulations, but we could qualitatively reproduce the picture from CD spectra when reweighting the ensembles against the *R*_h_ from experiments.

Furthermore, minor differences in the experimental conditions used in the SAXS and PFG NMR experiments^23^ might potentially lead to different behaviours of the ensembles in solution and hence some of the observed discrepancies. The Hst5 sample used for SAXS was in a buffer at pH 7.5 and ionic strength of about 150 mM, while the sample used for PFG NMR was at pH 7 and ionic strength of about 40 mM.^23^ Nonetheless, differences in ionic strength can affect the extent of inter-chain interactions.^45^ Furthermore, given that 7 out of the 24 residues of Hst5 are histidines, the difference in pH between the SAXS and NMR buffers can affect the protonation state of these residues and consequently affect the ensemble of conformations in solution. ^44^ To examine this, we performed coarse-grained simulations with the CALVADOS2 model^72,73^ using the IDRLab implementation on Google Colab.^74^ Although approximate, this model has previously been shown to capture some aspects of the ionic-strength dependency of chain compaction.^73^ The CALVADOS2 simulations of Hst5, however, suggest at most a small effect of pH and ionic strength (Table S1), though a more detailed analysis would ideally require experimental measurements or simulations with a better description of the effects of ions and mixed protonation states.^75^

In summary, we have here shown an example of the importance of using multiple and diverse experiments to obtain detailed insights into the conformational ensembles of IDPs, but also highlight some of the problems associated with analysing a diverse set of data. Even with such experiments in hand, we have also shown that it is important to have reliable models to calculate experimental observables from computational models. The development of these models is nonetheless impaired by the fact that it is far from trivial to generate accurate conformational ensembles of IDPs that can be used to train these models.^76^ Despite these caveats, we have used multiple experiments to shed further light on the structural basis of the temperature-dependent behaviour of Hst5. By generating conformational ensembles that reproduce the *R*_h_ from PFG NMR measurements, we support the hypothesis that there is a link between how temperature affects the amount of PPII structures and compaction.^15,23,77,78^ Some issues remain, that are related to whether the force field employed here underestimates the *R*_h_ of Hst5 or not. Future development of forward models, especially for CD, can help in better characterizing the content of PPII structures in IDPs. Furthermore, the study of more IDPs (ideally bigger, although much more difficult to sample efficiently by all-atom MD simulations) that show similar features would be useful to examine whether the behaviour observed for Hst5 occurs in other IDPs.

## Methods

### Molecular Dynamics Simulations

We performed MD simulations of Hst5 in solution with GROMACS 2018.6.^79^ We used the Amber99SB-disp force field^11^ with the TIP4P-D water model.^80^ The histidine residues in the peptide were modelled as their N*E* tautomers. We solvated the peptide in a dodecahedric box (with image distance equal to 10 nm) adding Na^+^ and Cl*^−^* ions to neutralize the system and reach an ionic strength of 150 mM (same as used in the experiments). We ran a two-steps energy minimization, using first the steepest-descent algorithm with energy tolerance set at 200 kJ mol*^−^*^1^ nm*^−^*^1^ and than the conjugate-gradient algorithm with energy tolerance set at 100 kJ mol*^−^*^1^ nm*^−^*^1^. Then, we equilibrated the system at the target temperature and pressure (1 bar) by running first a 200 ps MD simulation in NVT with position restraints on the protein atoms with the v-rescale thermostat,^81^ then a 1 ns MD simulation in NPT adding the Berendsen barostat.^82^ For production simulations, we used the Parrinello-Rahman barostat.^83^ We used a 2 fs time step in all simulations and constrained all bonds involving hydrogen atoms with the LINCS algorithm.^84^

To accelerate the sampling of the conformational space of Hst5, we employed Parallel Bias Metadynamics (PBMetaD)^56^ as enhanced sampling scheme. In PBMetaD a set of user-defined one-dimensional collective variables (CVs) is subject to history-dependent bias potentials that discourage the system from visiting regions of the CV space already explored. Biasing multiple one-dimensional CVs removes the limitation on the number of CVs that affects the classical metadynamics,^48^ where a single multidimensional space defined by the CVs is biased. Additionally, as in well-tempered metadynamics,^58^ the extent of biasing decreases when the same region of a CV space is visited multiple times, so to have a smoother convergence than with classical metadynamics, and the free-energy surface (FES) as a function of the CVs can be estimated within a well-defined error.^85^ In our PBMetaD setup we used 47 CVs, consisting in all 23 *φ* and all 23 *ψ* dihedral angles and the radius of gyration of the C*_α_* atoms (*R*_g_*_,Cα_*). We employ *R*_g_*_,Cα_* as CV to bias because it is cheaper to compute on-the-fly, but in all further analyses we calculate the *R*_g_ for all the atoms. Gaussian bias was deposited every 400 ps. We set the initial height of the Gaussian bias to 0.3 kJ mol*^−^*^1^. This height decreased during the simulations by means of the well-tempered feature, according to a bias factor of 24. We used an adaptive algorithm to define the width of the Gaussian bias potentials,^86^ where, at each bias deposition, the width is based on the space covered by the CV in the last 2 ps. To enhance the sampling efficiency even further we used a double walker approach,^57^ where two simulations run in parallel sharing the deposited bias. For the CV representing the *R*_g_*_,Cα_*, we set an interval in the CV space between 0.6 and 2.5 nm outside which the biasing force of the PBMetaD does not act. ^87^ We did so because we observed a ‘FES wall climbing effect’: since the FES as a function of the *R*_g_*_,Cα_* is shaped as a single basin, as bias accumulates, new areas in the CV space, corresponding to more compact and more extended high-energy conformations, gets available (Fig. S11). Since these areas are newly explored, bias is deposited with the initial height, while for the rest of the CV space the extent of biasing is already very low. The advantage of the well-tempered feature in PBMetaD is that when all the accessible conformations in CV space are explored, the biasing potential decreases at the point of becoming quasi-static and the simulation converges to a well definite FES. But if new areas of the CV space become continuously accessible and a large bias is deposited, the convergence is impaired. Setting the interval in the FES of the *R*_g_*_,Cα_*, prevents the system from climbing the FES walls corresponding to high-energy conformations that are not relevant. Despite the use of the interval in the FES of the *R*_g_, we still observe few frames depositing a large amount of bias outside the interval (Fig. S4). This, however, is not a problem since our calculations are not based on those few frames that explore regions of the FES outside the interval range. To enable PBMetaD in GROMACS we used PLUMED 2.5.^88^

At the end of a simulation, we recovered the total bias (*B*) for each conformation *x* explored and used it to calculate the Boltzmann weights (*ω*) associated to each conformation according to 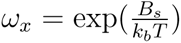, where *k_b_* is the Boltzmann constant and *T* is the temperature. The FES for a CV is calculated from the weighted histogram (*H*) of the CV as *FES*(*CV*) = *−k_b_T* ln *H*(*CV*). The error on the FES and ensemble averaged values is calculated by block averaging.^85,89^ The package used for these calculations is available at https://github.com/fpesceKU/BLOCKING.

### Calculation of experimental observables

We compare the ensembles from our simulations to SAXS, PFG NMR and CD data. We calculate SAXS data with Pepsi-SAXS,^90^ following the protocol we previously described:^69^ the intensity of the forward scattering and constant background are fit to the experimental SAXS profile by least square fitting; the values for the density of the hydration layer and the excluded volume are scanned on a two-dimensional grid and the combination of those that best fit the data upon reweighting is selected (Table S2).

We calculate the *R*_h_ using the HullRadSAS software^64^ for reasons recently described.^62,63^ We calculate the per-residue secondary structure with DSSP-PPII ^91^ where—if two or more consecutive residues that are not assigned to any secondary structure by the DSSP algorithm^92^ have backbone dihedrals correspondent to those typical of PPII structure—those residues are assigned as PPII structure. For each residue we then report the propensity for PPII structure formation as a weighted sum of the frames where a residue forms PPII structures, using either weights from PBMetaD or BME (see below).

### Experimental data

All experimental data used in this study were previously measured and published by Jephthah et al.^23^ We corrected the published *R*_h_ values based on a recent estimate of the reference *R*_h_ value for 1,4-dioxane (2.27 Å *±* 0.04 Å^63^ compared to the previously used 2.12 Å^65^). Furthermore, we rescaled the error bars for the SAXS intensities by a factor estimated through the Bayesian indirect Fourier transform.^93^

### Ensemble reweighting

We use the Bayesian/Maximum entropy (BME) approach^53^ to reweight simulations in order to improve the agreement with experimental data. This is achieved by minimizing the functional *L*:^52,54^

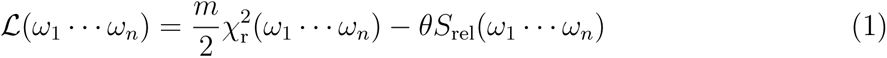

where *m* is the number of experimental data points, (*ω*_1_ *…ω_n_*) are the weights for each simulation frame, *χ*^2^_*r*_ measures the agreement between the calculated and experimental observable. The relative entropy, 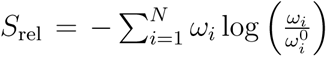, indicates how much the refined weights (*i.e.* the posterior distribution, *ω*) diverge from the initial weights (*i.e.* the prior distribution, *ω*^0^), *θ* is a free-parameter that balances the extent of minimizing the *χ*^2^_*r*_ vs. maximizing *S*_rel_. By taking the exponential of *S*_rel_ (*φ*_eff_ = exp(*S*_rel_)), it is possible to gain insights into the extent of the reweighting. Generally, a low *φ*_eff_ is not desirable, as it means that a low number of frames from the simulation contributes to the calculated weighted averages. Therefore different values for *θ* are explored in the reweighting procedure, and that corresponding to the *χ*^2^_*r*_ reaching plateau with the highest *φ*_eff_ is chosen to generate the reweighted ensemble.^53^ This helps to ensure a posterior distribution with a good agreement with the experimental data while retaining as much information as possible from the prior distribution. The prior distribution, *ω*^0^ is estimated as the weights from PBMetaD.

To reweight simulations against *R*_h_ values from PFG NMR we use the standard implementation of BME. For SAXS data we use the iterative BME (iBME).^69^ iBME couples ensemble reweighting with least square fitting of scale factor (*s*) and constant background (*cst*) to the calculated SAXS profile. Therefore here the *χ*^2^_*r*_ in the *L* functional takes the following shape:

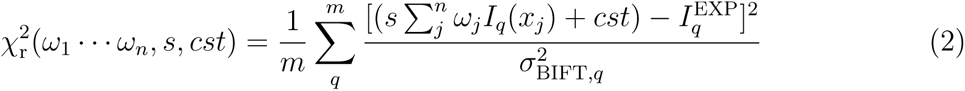

where *I_q_*(*x_j_*) is the calculated SAXS intensity at scattering angle *q* for the frame *x_j_*, *I*^EXP^_*q*_ is the experimental SAXS intensity at scattering angle *q*. *σ*_BIFT_*_,q_* is the rescaled error^93^ of the experimental intensity at scattering angle *q*.

In reweighting simultaneously against *R*_h_ and SAXS we also use the standard BME, using calculated SAXS profiles with scale factor and offset optimized from previous iBME runs.

## Data and code availability

MD simulations are available at https://doi.org/10.17894/UCPH.4D0D9A09-B40B-4A3A-93EF-D6C249F49A90. PLUMED inputs to enable PBMetaD are available on PLUMEDNEST^94^ at https://www.plumed-nest.org/eggs/23/008/. Scripts used to analyze the simulations are available at https://github.com/KULL-Centre/_2023_Pesce_Hst5.

## Acknowledgement

We thank Marie Skepö and Birthe B. Kragelund for sharing the SAXS, NMR and CD data and for discussions on disordered proteins, and Simone Orioli for discussions about the simulations. This research was funded by the Lundbeck Foundation BRAINSTRUC initiative in structural biology (R155-2015-2666, lundbeckfonden.com). We acknowledge access to computational resources from the ROBUST Resource for Biomolecular Simulations (supported by the Novo Nordisk Foundation grant no. NF18OC0032608) and from the Biocomputing Core Facility at the Department of Biology, University of Copenhagen.

