## Supporting Figures and Tables for "Combining experiments and simulations to examine the temperature-dependent behaviour of a disordered protein"

### Supplementary material for: Combining experiments and simulations to examine the temperature-dependent behaviour of flexible peptides

Francesco Pesce and Kresten Lindorff-Larsen*

*Structural Biology and NMR Laboratory, The Linderstrøm-Lang Centre for Protein Science, Department of Biology, University of Copenhagen, Copenhagen, Denmark*

#### Supplementary figures and tables

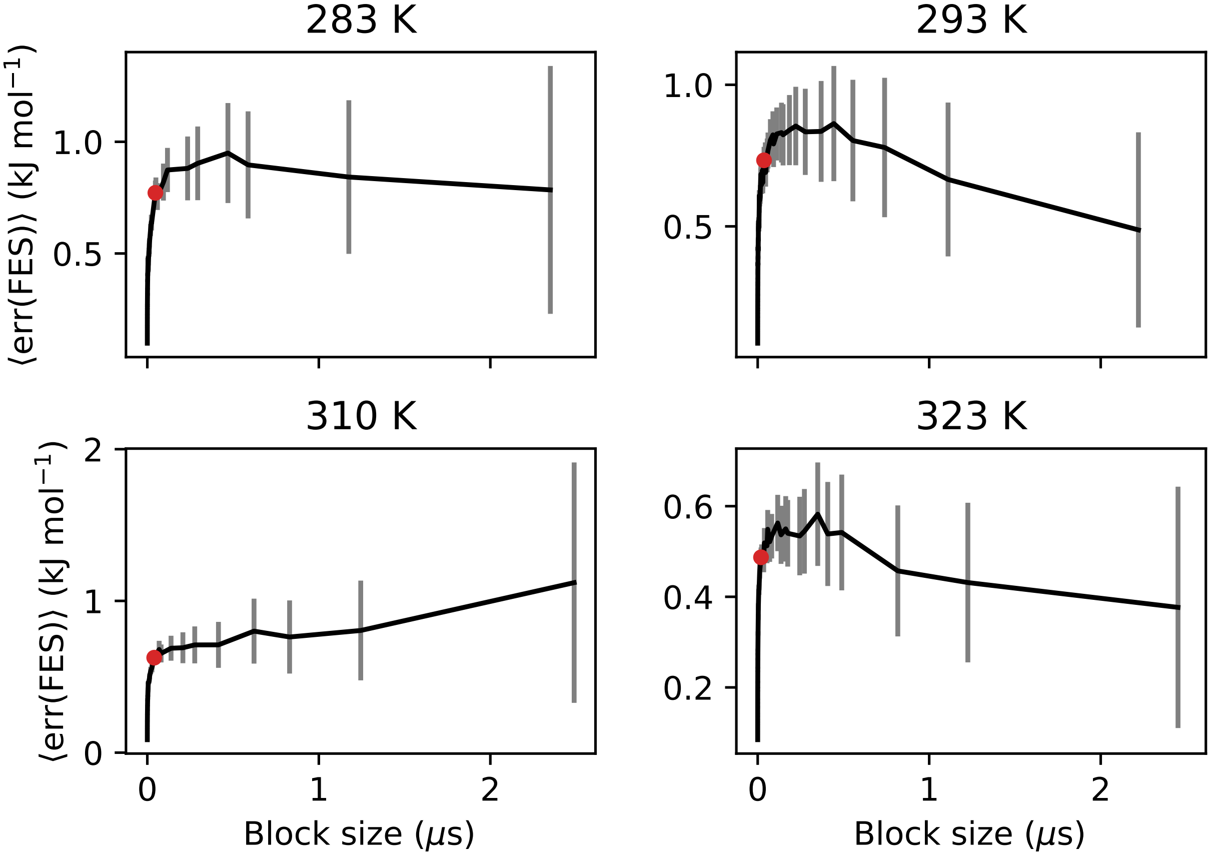


Figure S1: (a) Block analysis on the free-energy surface as a function of the $R_\mathrm{g}$. As in the more common block analysis applied to correlated data, we divide the simulations in blocks. But instead of looking at the standard error of the averages over the blocks, we calculate the free-energy surface for each block and calculate the standard error of the free-energy over the blocks. Each point in the block then represent the average standard error over the free-energy bins for a certain block size. We note that we here perform the analysis over both replicas from our multiple-walker simulations, and so the maximal block size corresponds to the length of each of these simulations (corresponding to two blocks). The red dot at the beginning of the plateau region of the curves represent the block size used for the calculation of error bars of $R_\mathrm{g}$ and $R_\mathrm{h}$.

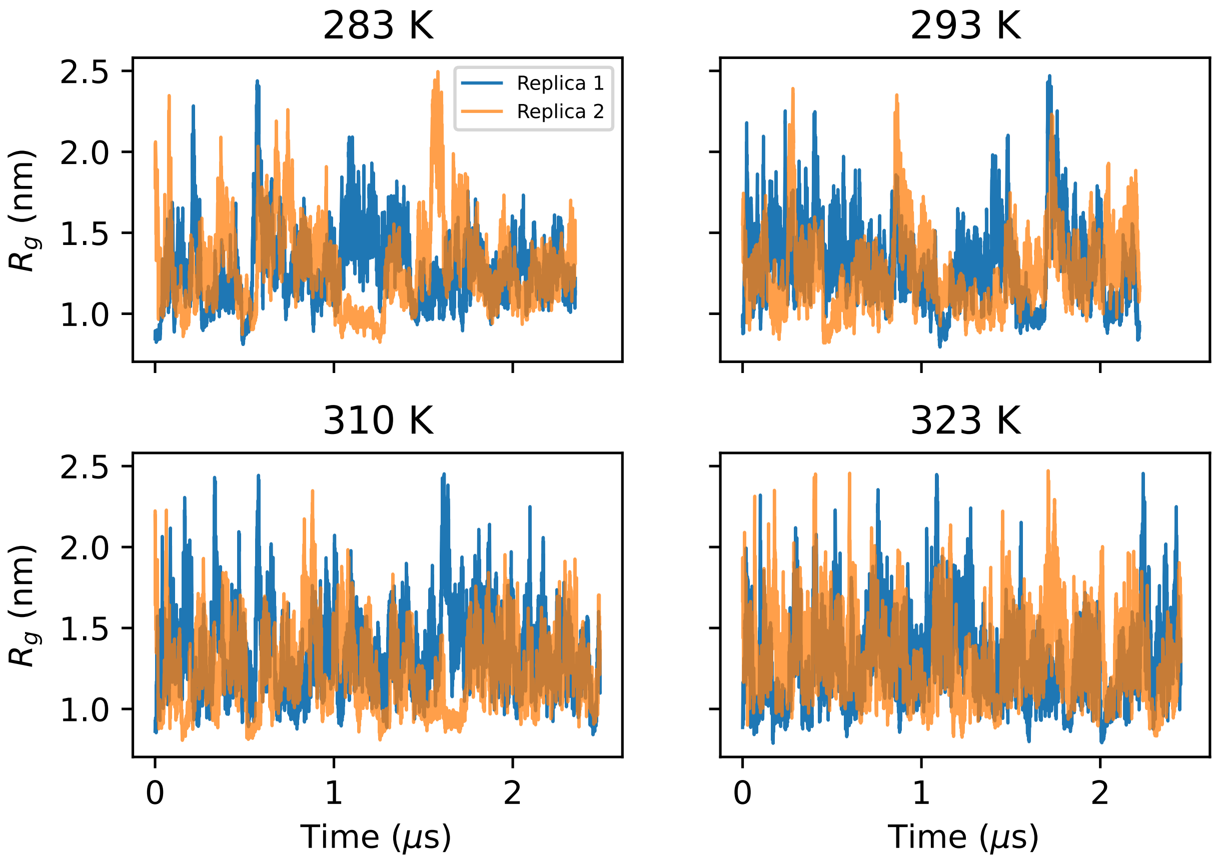


Figure S2: Sampling of the $R_g$ in the PBMetaD simulations with two replicas.

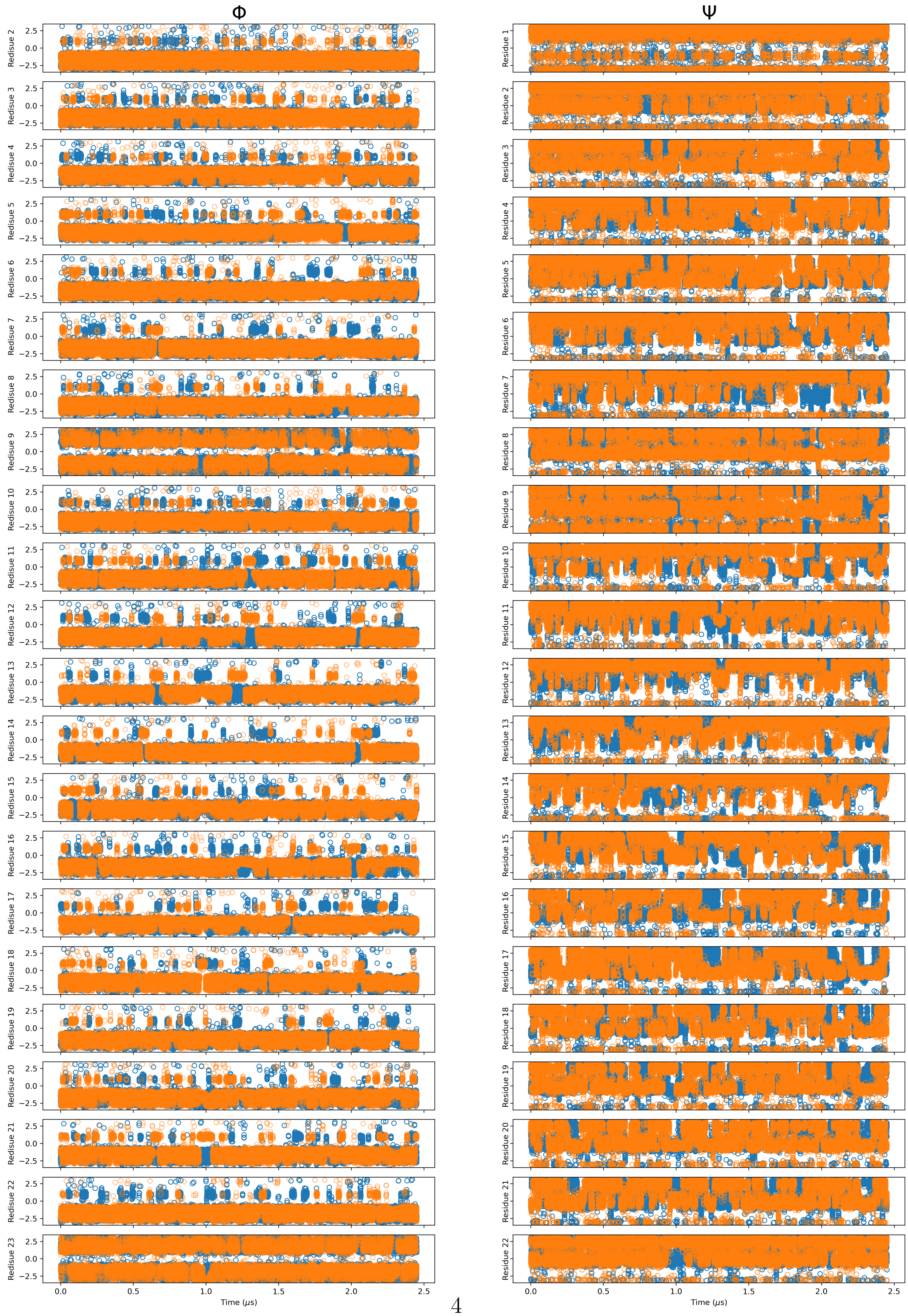




Figure S3: Sampling of the backbone dihedrals in the PBMetaD simulations with two replicas. Units on the y-axes are in radians.

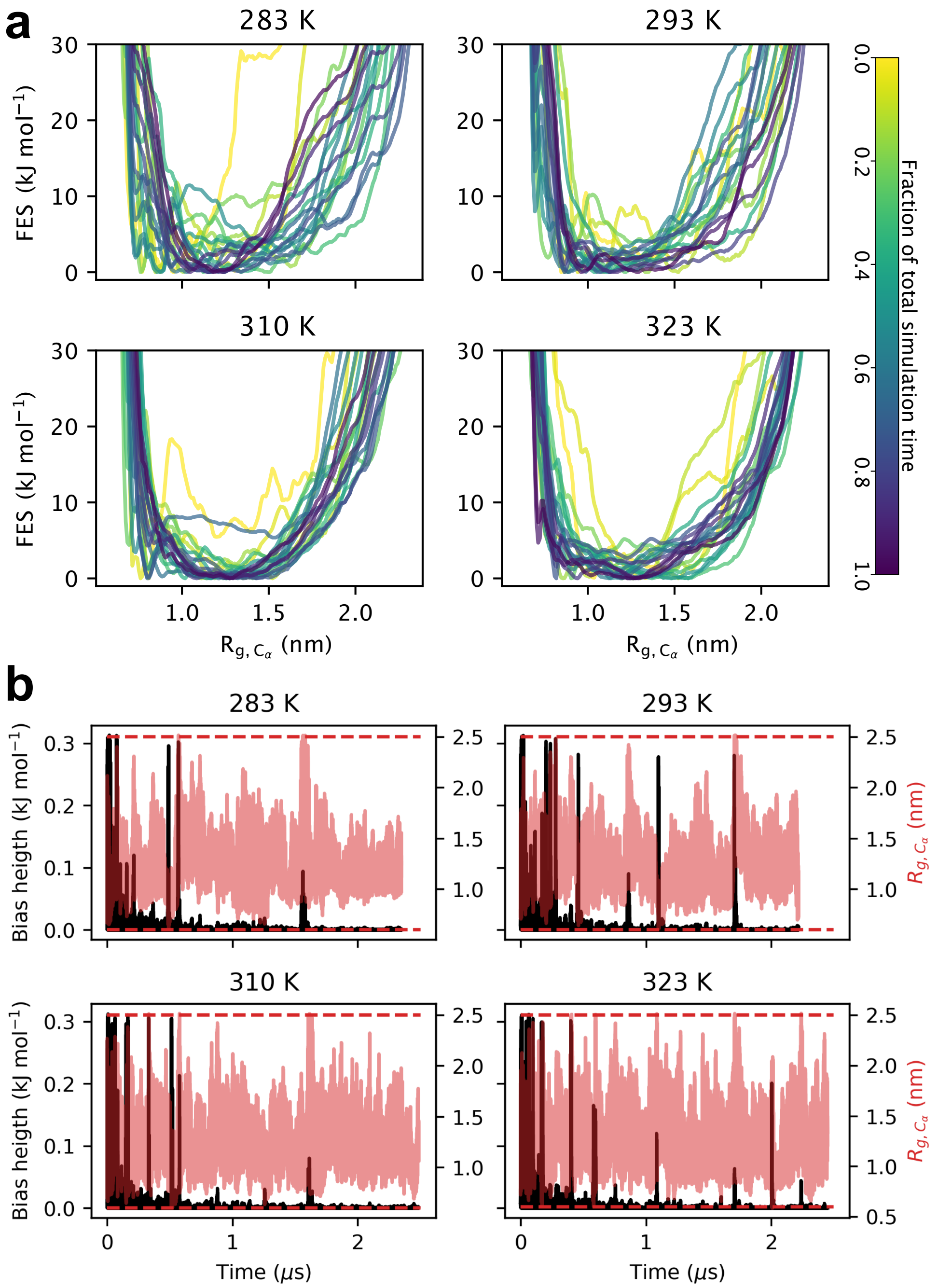


Figure S4: (a) Time-series of the bias deposition (black) on the $R_{g,C_\alpha}$ collective variable for two replicas (red) in the PBMetaD simulations. The horizontal dashed lines in red mark the interval outside of which the system does not experience the biasing force (lower interval at 0.6 nm and upper interval at 2.5 nm). (b) FES of the $R_g$ reconstructed from the bias accumulated is shown at different fractions of the total simulation time. Because of the well-tempered metadynamics, the amount of deposited bias decreases over time.

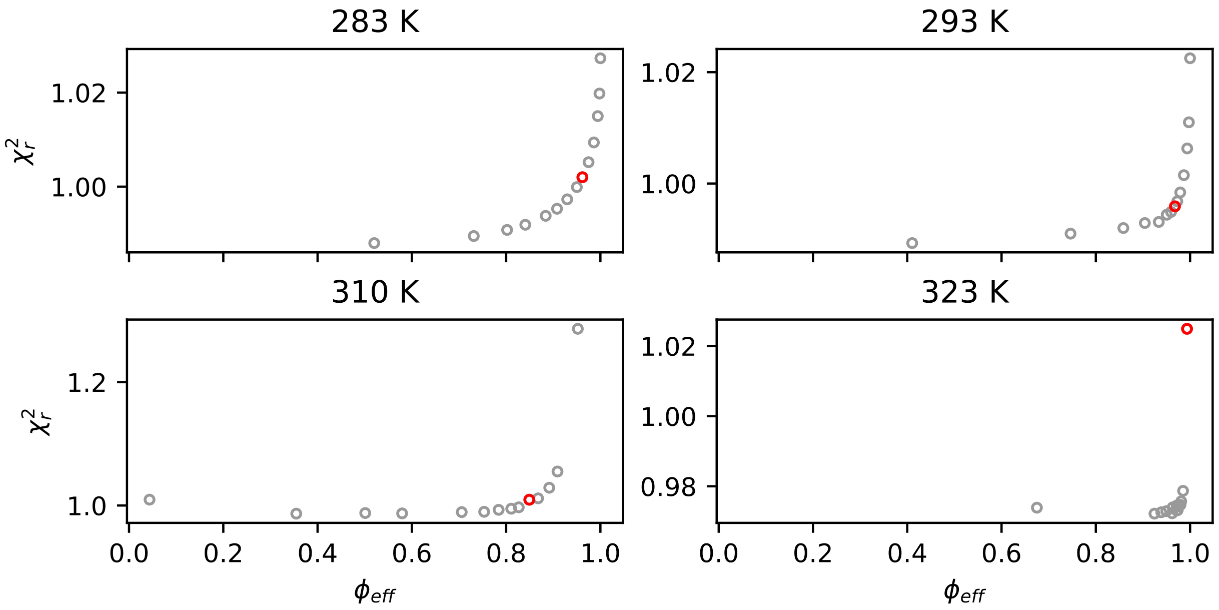


Figure S5: Scan of $\theta$ values used in reweighting ensembles against SAXS data with the iBME approach. For each ensemble, the $\theta$ corresponding to the red marker was used (100, 75, 150 and 10000 respectively for simulations and 283 K, 293 K, 310 K and 323 K).

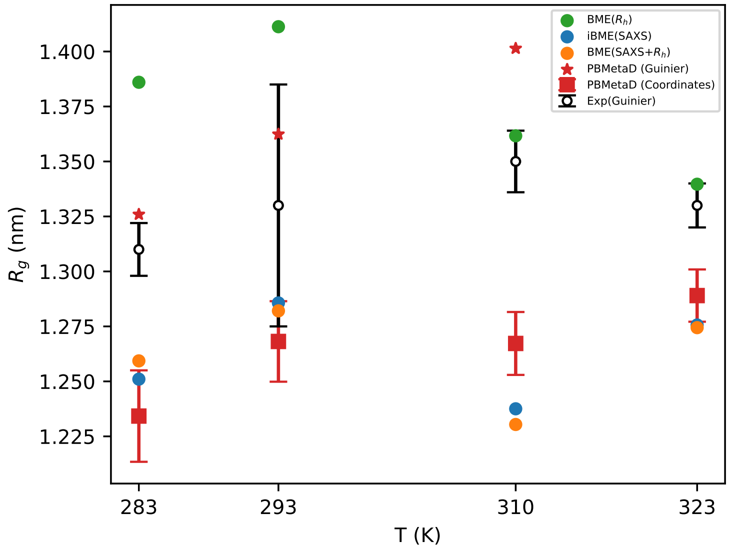


Figure S6: Ensemble-averaged $R_g$ from the simulations at different temperatures (red), calculated from atomic coordinates (square markers) or from Guinier fitting of the calculated SAXS profile (star markers), reweighted against SAXS (blue), $R_h$ (green) or both $R_h$ and SAXS (orange). The $R_g$ from Guinier fitting of the experimental SAXS profile is shown in black.

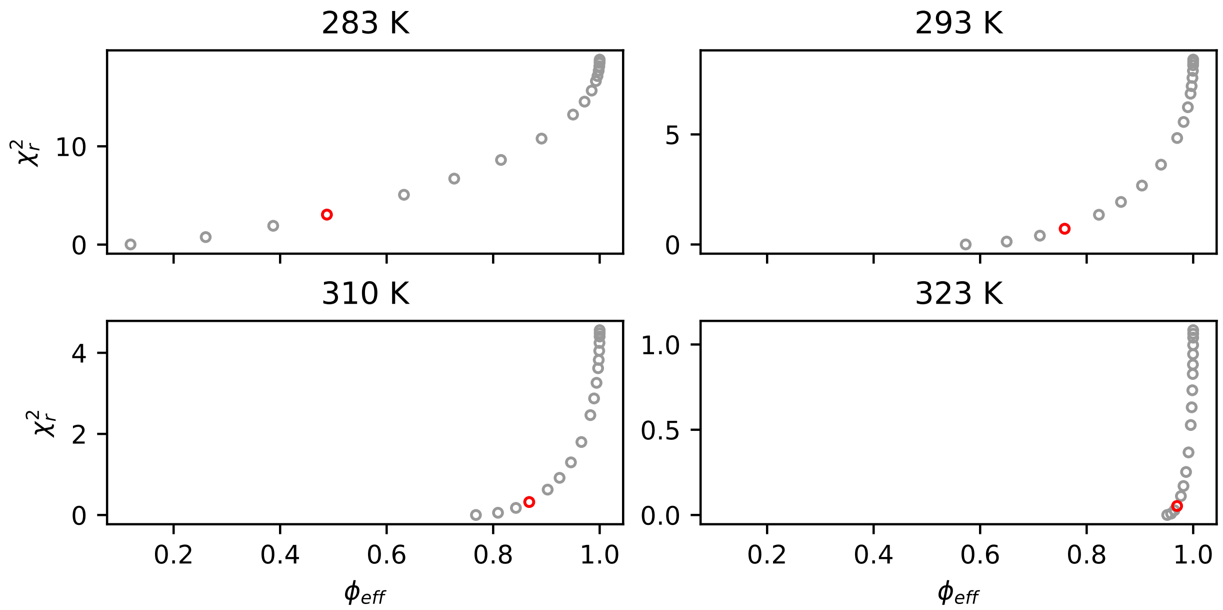


Figure S7: Scan of $\theta$ values used in reweighting ensembles against $R_h$ from PFG NMR with the BME approach. For each ensemble, the $\theta$ corresponding to the red marker was used ($\theta$=3 in all cases).

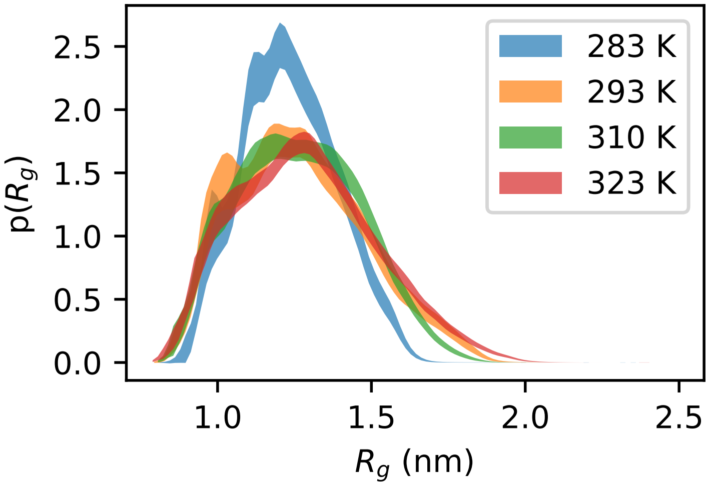


Figure S8: Distributions of $R_g$ from PBMetaD simulations at different temperatures, including errors on the probability estimate from block analysis.

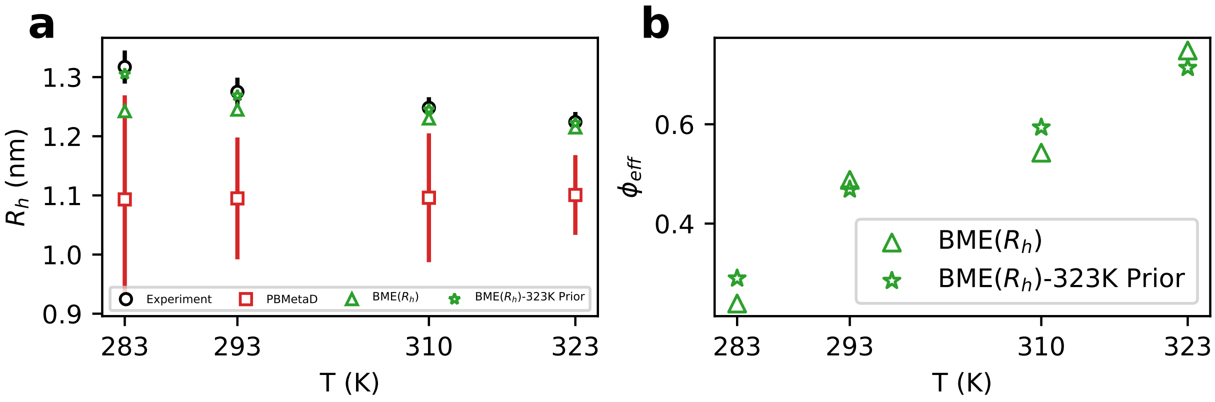


Figure S9: Reweighting simulations at different temperatures against $R_\mathrm{h}$ from experiments at the respective temperatures (green triangle) or reweighting the simulation at 323 K against experimental $R_\mathrm{h}$ at different temperatures (green star). (a) comparison of the resulting reweighted $R_\mathrm{h}$ with that from simulations (red squares) and the experiments (black circles). (b) effective fraction of frames contributing to reweighted averages ($\phi_\mathrm{eff}$) when reweighting all simulations (with $\theta$=5) or that at 323 K (with $\theta$=1).

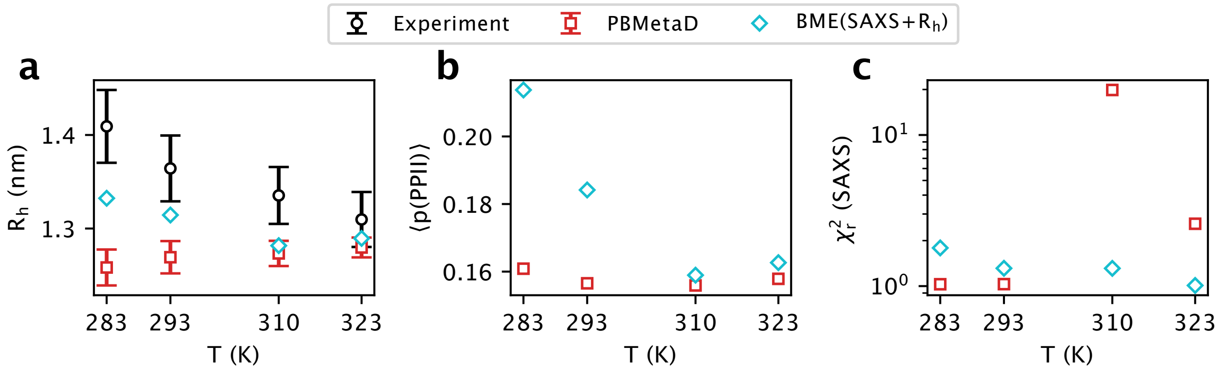


Figure S10: Optimization of the simulations against SAXS and $R_\mathrm{h}$ simultaneously. In contrast to Fig. 4 in the main text, we here use the same number of data points for SAXS and $R_\mathrm{h}$. We show the (a) $\chi_r^2$ to SAXS data (b) $R_\mathrm{h}$ and (c) average PPII structure propensity from the experiment (black, when applicable), the ensemble average from the simulations (red) and the simulations reweighted against both SAXS and $R_\mathrm{h}$ (cyan).

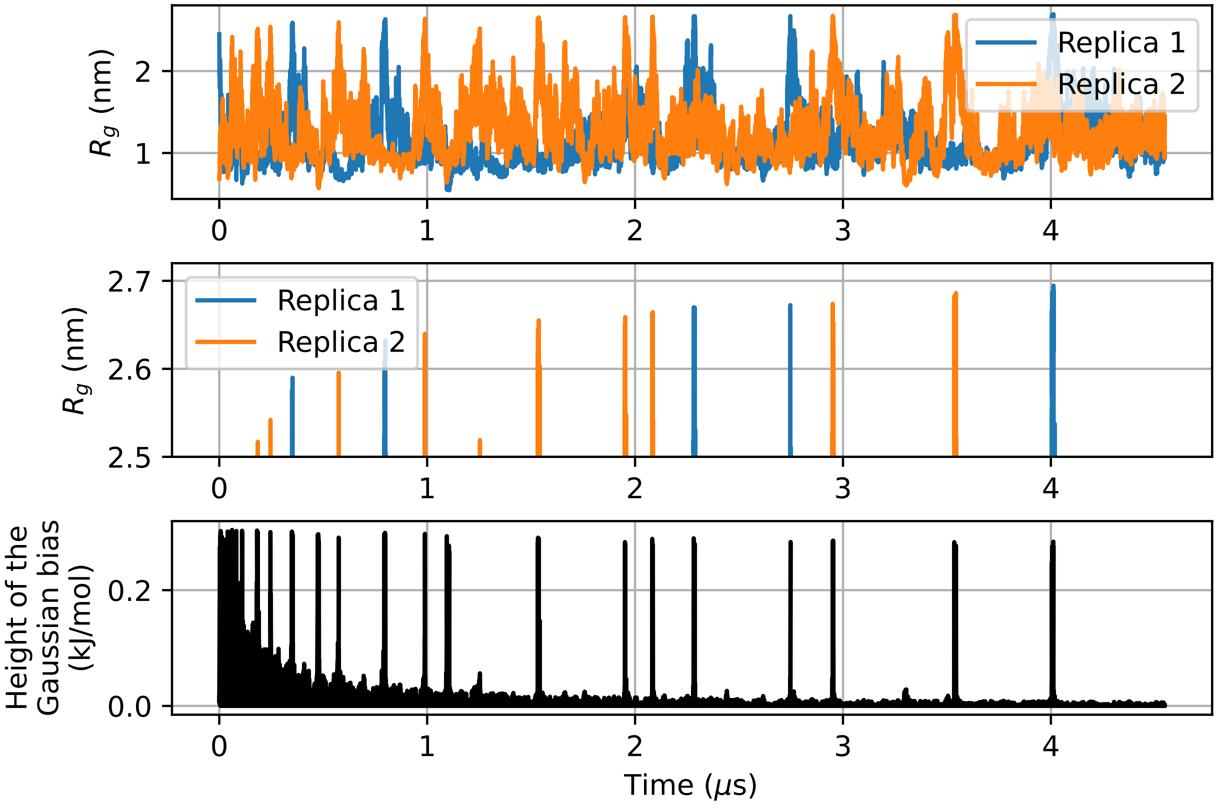


Figure S11: PBMetaD simulation at 310 K not employing intervals in biasing the $R_{\mathrm{g}}$. We show how the system keeps reaching more and more extended conformations, and the corresponding deposition of a large amount of bias, impairing the convergence of the FES. Top panel: full time series of the $R_{\mathrm{g}}$ in the simulation. Middle panel: zoom on the high-$R_{\mathrm{g}}$ regions explored. Bottom panel: bias deposition along the simulation.

Table S1: $R_g$ values (in nm) calculated from CALVADOS2 coarse-grained simulations at different values of pH and ionic strength (IS).

| | | **Ionic strength (mM)** | |
|---|---|---|---|
| | | **40** | **150** |
| **pH** | **7.0** | 1.290 ± 0.008 | 1.278 ± 0.008 |
| | **7.5** | 1.287± 0.008 | 1.282 ± 0.008 |

Table S2: Pepsi-SAXS parameters for the density of the hydration layer ($\delta\rho$) and excluded volume ($r\_0$) that resulted in the SAXS-optimized ensembles with lowest $\chi_r^2$ and highest $\phi_{\text{eff}}$.

| | 283 K | 293 K | 310 K | 323 K |
|---|---|---|---|---|
| $\delta\rho$ | 8.000 | 8.000 | 6.000 | 6.00 |
| $r\_0$ | 1.025 | 1.025 | 1.075 | 1.05 |
